## Supplementary information for "Deep mutational scanning of essential bacterial proteins can guide antibiotic development"

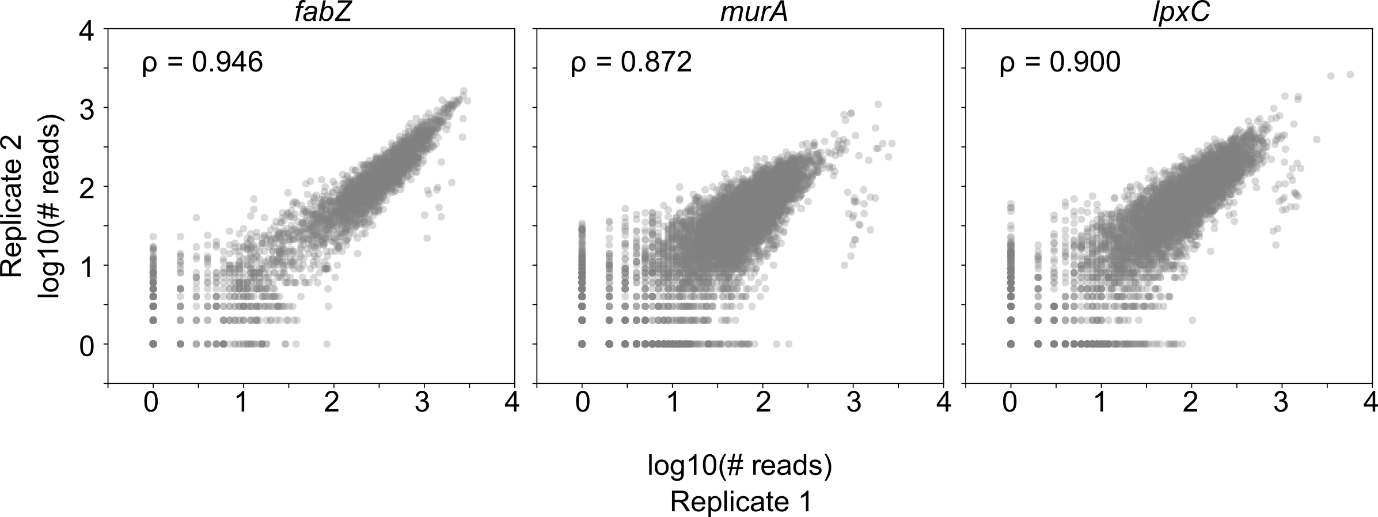

**Figure S1: Library construction by Onyx® is highly reproducible.** The number of Onyx® edit reads detected per edit across two independent library builds on the Onyx® instrument for each gene library is shown. Spearman’s rank correlation coefficients are indicated, which demonstrate the high reproducibility in library construction.

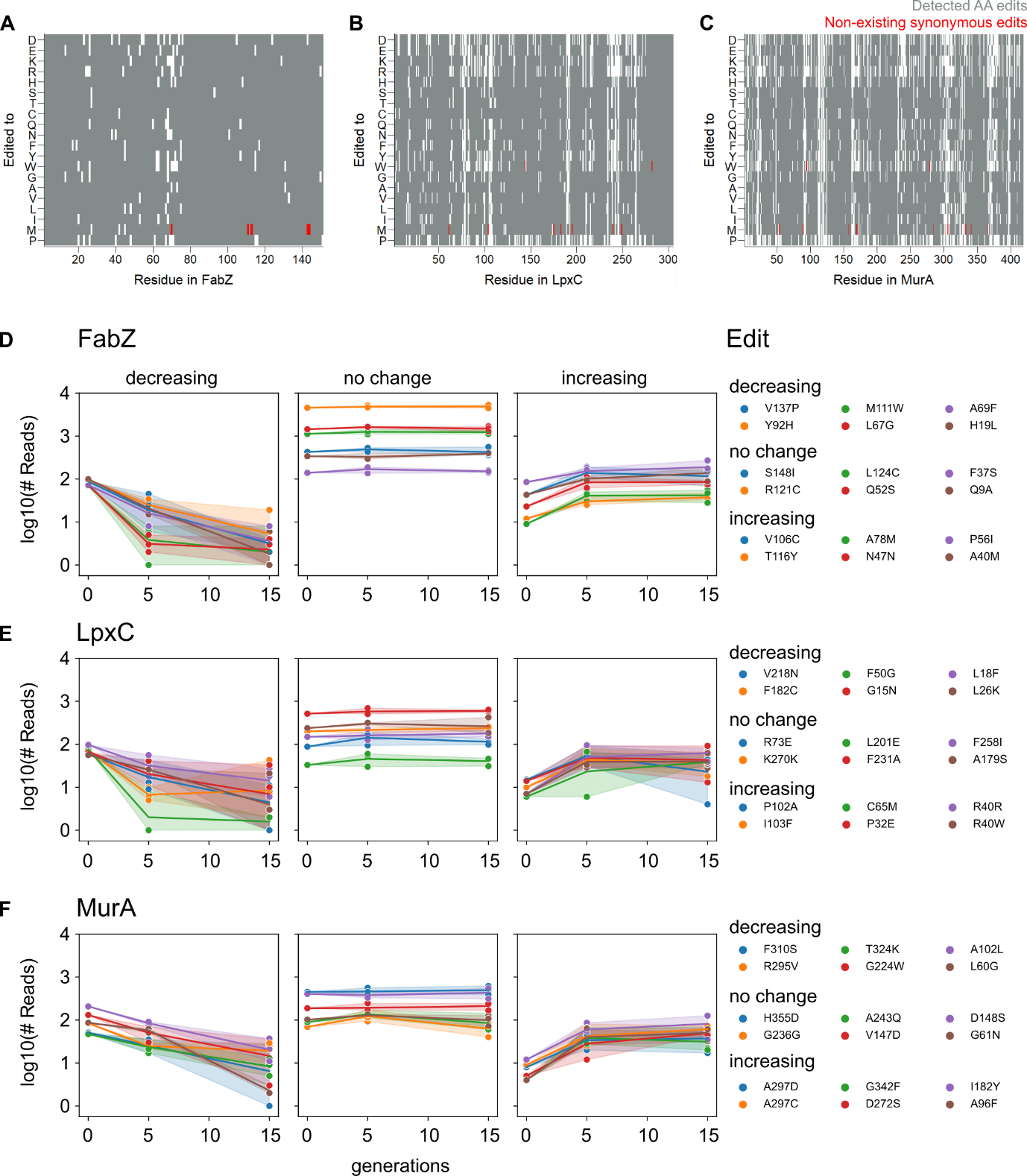

**Figure S2: Edit read counts determined right after construction of saturation mutagenesis libraries reveal biologically relevant information but continue to change throughout early growth cycles.** A-C) Heat maps indicating the presence or absence of specific amino acid substitutions in the saturation editing library of FabZ (A), LpxC (B) and MurA (C). Edits that were not detected in the library are shown in white, while detected amino acid substitutions are colored grey. Those synonymous edits that cannot be designed due to the absence of a synonymous codon (M and W) are shown in red. D-F) Libraries were grown and sequenced to determine library composition after 0, 5 and 15 generations. Read counts were plotted for a number of randomly selected edits from the FabZ (D), LpxC (E) or MurA (F) libraries that either show no change in read counts over time or have increasing or decreasing read counts.

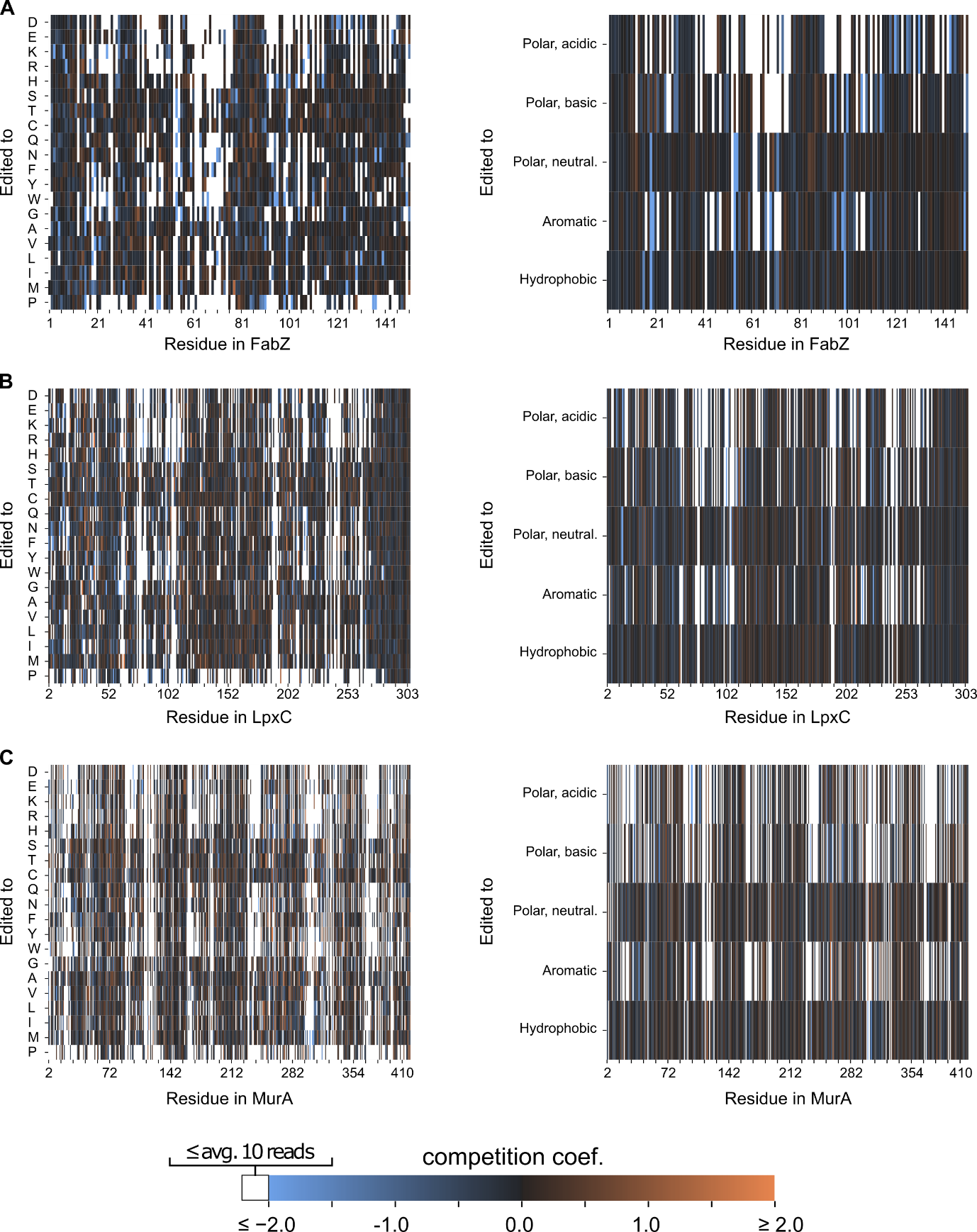

**Figure S3: Competition within saturation editing libraries identifies fitness effects of detected edits.** To interrogate the fitness effects associated with all detected edits, libraries were grown to allow for competition between variants. After 0, 5 and 15 generations of growth, library composition was determined by Illumina sequencing. The change in abundance of each variant was fitted and the normalized and adjusted slope of the fit was taken to be a mutation’s competition coefficient (see Materials & Methods). Heat maps of competition coefficients associated with the FabZ (A), MurA (B) and LpxC (C) libraries are shown, where edits are shown individually (left) or grouped based on the biochemical properties of the introduced amino acid (right).

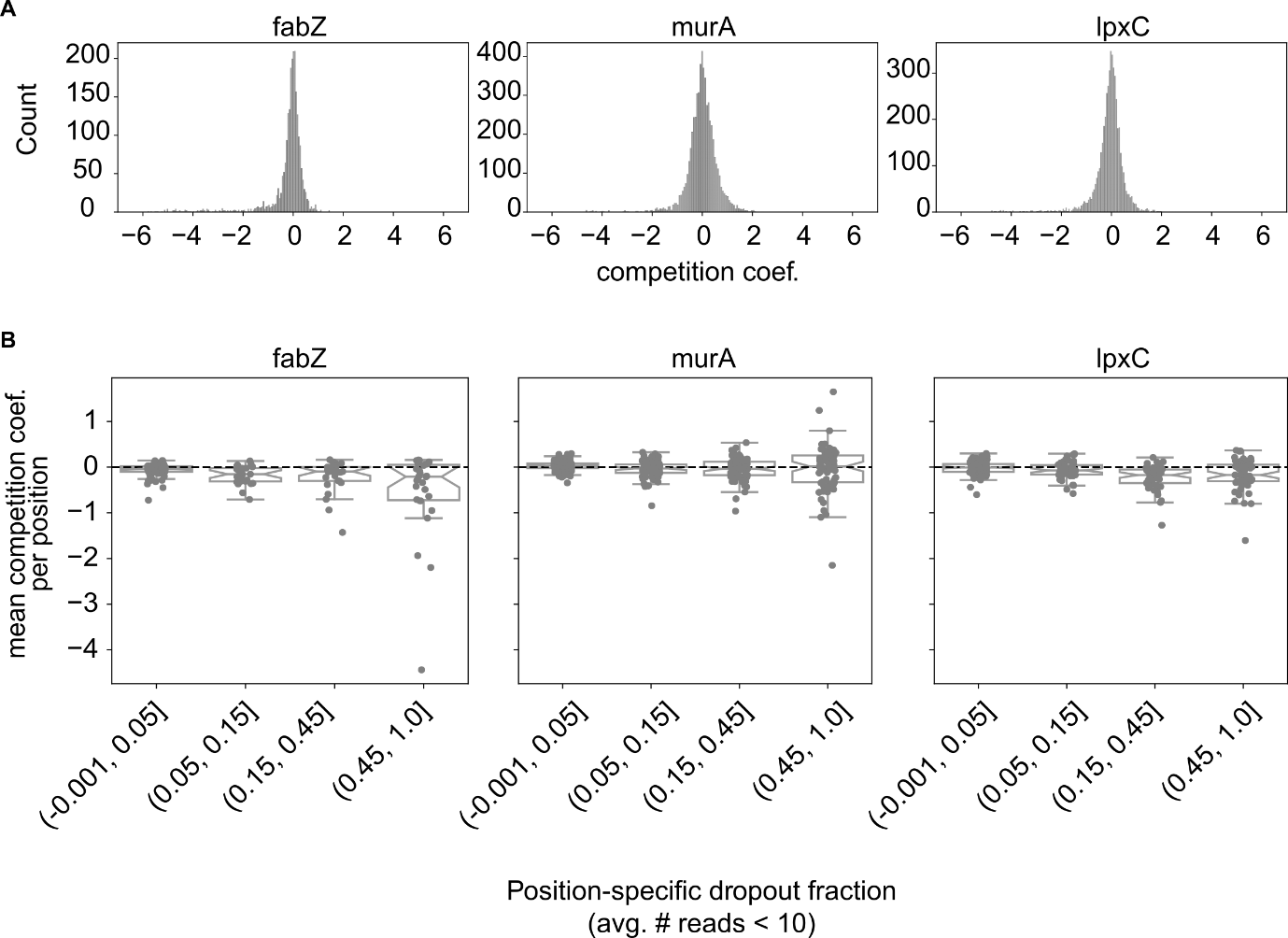

**Figure S4: Competition coefficients can be used to characterize fitness effects of individual edits.**

To interrogate the fitness effects associated with all detected edits, libraries were grown to allow for competition between variants. After 0, 5 and 15 generations of growth, library composition was determined by Illumina sequencing. The change in abundance of each variant was fitted and the normalized and adjusted slope of the fit was taken to be a mutation’s competition coefficient (see Materials & Methods). A) The distribution of competition coefficients for each library is shown. B) By plotting the mean competition coefficient grouped by the dropout fraction of the position in question, it becomes clear that residues associated with higher dropout fractions also have more negative competition coefficients, meaning that amino acid changes at these positions are generally more harmful and deleterious to protein function.

**Table S7: Selected *murA* mutations are not causal for fosfomycin resistance.** Selected *murA* mutant alleles were transferred to a clean genetic background and fosfomycin MIC values of the wild-type parent strain and the *murA* mutants were determined. Relative MIC values were determined compared to wt. Wt, wildtype.

| **Strain** | **Fosfomycin MIC (µg/ml)** | **Relative MIC** |
| --- | --- | --- |
| wt | 8 |  |
| MurA V16M | 8 | 1x |
| MurA D51A | 4 | 0.5x |
| MurA V228I | 4 | 0.5x |
| MurA R267E | 8 | 1x |
| MurA I402W | 4 | 0.5x |
